# How AlphaFold shaped the structural coverage of the human transmembrane proteome

**DOI:** 10.1101/2023.04.18.537193

**Authors:** Márton A. Jambrich, Gabor E. Tusnady, Laszlo Dobson

## Abstract

AlphaFold2 (AF2) provides structures for every protein, opening up prospects for virtually every field in structural biology. However, transmembrane proteins pose a challenge for experimental scientists, resulting in a limited number of solved structures. Consequently, algorithms trained on this finite training set also face difficulties. To address this issue, we recently launched the TmAlphaFold database, where predicted AlphaFold2 structures are embedded into the membrane and a quality assessment is provided for each prediction using geometrical evaluation. In this paper, we analyze how AF2 has changed the structural coverage of membrane proteins compared to earlier years when only experimental structures were available, and high-throughput structure prediction was greatly limited. We also evaluate how AF2 can be used to search for (distant) homologs in highly diverse protein families. By combining quality assessment and homology search, we can pinpoint protein families where AF2 accuracy is limited, and experimental approaches are still desired.

## Introduction

Transmembrane (TM) proteins are gatekeepers that regulate the movement of nutrients and drugs in and out of cells while also filtering or amplifying signals in neurotransmission and perception. After the Human Genome Project, it was determined that around 20-30% of the coded protein in the human genome encodes transmembrane proteins (TMPs), which means that there are approximately 5-7 thousand TMPs in the human proteome^1^. The lipid bilayer surrounding the TMPs separates them into distinct phases, which require a multi-step folding process to properly encapsulate them^2^. Despite the crucial role that TMPs play in cellular processes, their structural characterization lags far behind that of globular proteins due to their natural dual environment^3^, which makes their purification and crystallization difficult^4^.

The challenging experimental conditions have given rise to various computational approaches, ranging from topology prediction^5^ (which determines the exact location of TM segments and the orientation of connecting loops relative to the membrane, e.g., cytosolic, extracellular), to 3D prediction of TMPs^6^. The latter has become a popular field, with approaches borrowed from the prediction of globular proteins and extended with special conditions arising from the topology during contact map definition^7^. Despite enormous efforts, the accuracy of these topology and 3D predictions is limited, because the training set for tuning these methods is limited by the number of experimentally solved structures. In the 2000s numerous structural genomic target selection projects began to pave the way for membrane protein structure determination^8,9^, with limited success. The situation has improved over the last 10 years, in particular as cryo-electron microscopy (cryo-EM) has revealed hundreds of new TMP structures.

But the breakthrough seems to come from a computational tool: AlphaFold2^10^ (AF2), developed by the team at DeepMind, has revolutionized the field of structural biology by greatly improving the speed and accuracy of protein structure prediction. The software uses deep learning algorithms and a vast amount of protein sequential and structural data to predict the 3D structure of a protein based solely on its amino acid sequence. It was quickly realized that AF2 could be used in many ways for structural biology research, including experimental modelling, prediction of intrinsically disordered regions, or assessment of disease-causing mutations^11^. However, the question remained as to how AF2 performs in the field of membrane proteins where the training data it could use was still very limited.

We have recently launched the TmAlphaFold database^12^, which adds the possible localization of the lipid bilayer relative to the AF2-predicted helical TMP structures and evaluates the quality of each prediction. In this study, we analyze the human TM proteome to identify bottlenecks where we encountered limitations with the AF2 structures. We made a quality assessment of the state of the human TM proteome, performed sequence- and structure-based clustering of TMPs, and then highlighted proteins or protein families where there is still room for improvement.

## Results

### TmAlphaFold database is a good starting point to discriminate good and bad quality structures

The TmAlphaFold database^12^ not only provides a possible localization of the lipid bilayer in the predicted structures but also evaluates the quality of the predicted membrane-embedded structures by applying filters, such as globular domains embedded in the membrane, conflicts with the topology prediction, highlighting membrane regions folded outside the bilayer, and more (see TmalphaFold web page for details). We constructed a non-redundant human TMP benchmark set consisting of proteins with I) solved 3D structures^3^ and II) high quality topology predictions from the Human Transmembrane Proteome (HTP) database^1^. First, we checked whether the different filters capture unique errors to rule out that they all detect the same problems but with a different approach (Figure 1A). Using the benchmark set, we calculated the correlation between the filters and found that most of the filters are diverse.

**Figure 1.**
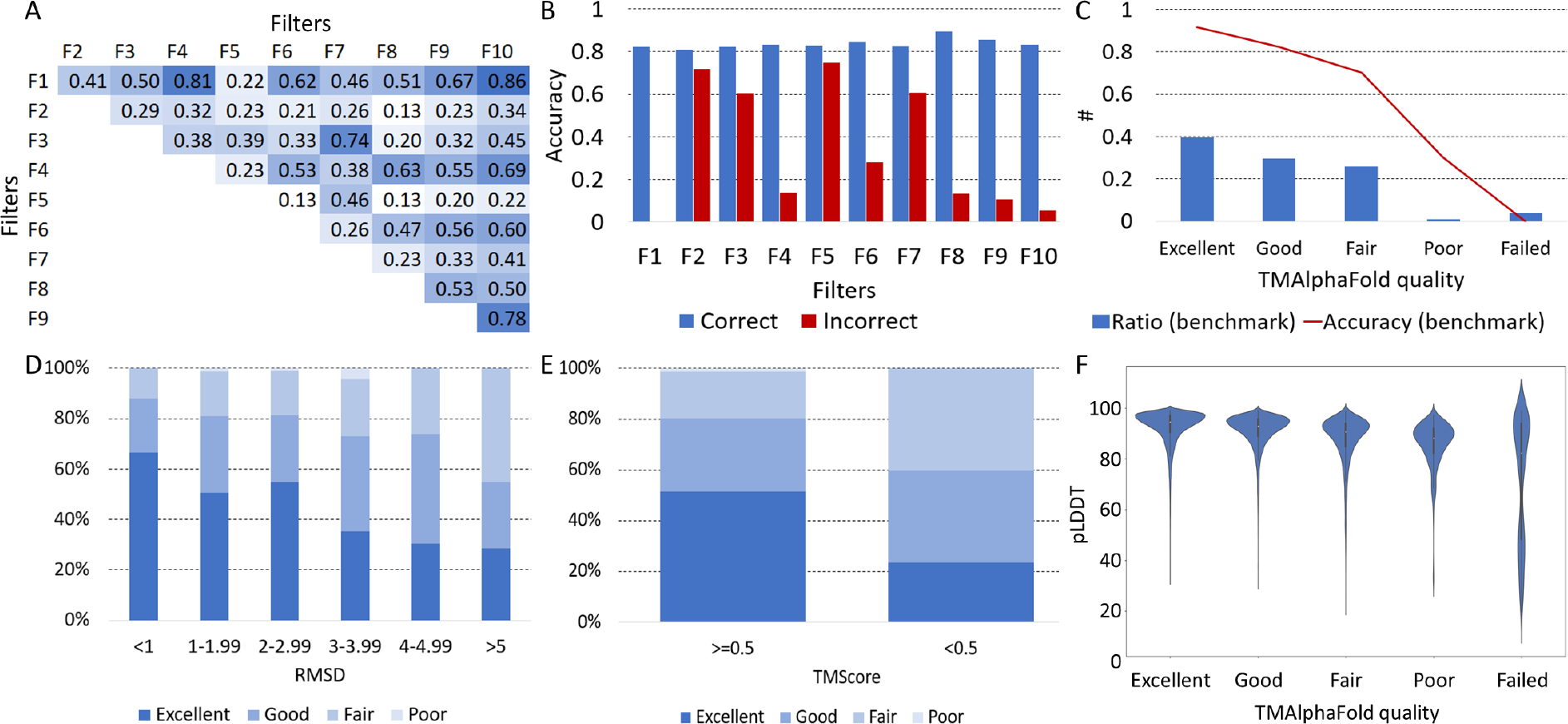
A) Correlation of different filters from TMAlphaFold on the benchmark set (F1: DetectingMembranePlane, F2: Signal, F3: FullStructure, F4: ShortHelix, F5: Masked, F6: MissingTmpart, F7: Domain, F8: OverpredictCctop, F9: UnderpredictCctop, F10: MembranePlaneCctop). Darker shade of blue means higher correlation; B) Performance of different filters on correct (topographically correct) and incorrect structures in the benchmark set; C) Ratio of proteins and the measured accuracy (by checking topography) in different quality levels on the benchmark set; D) Distribution of TMAlphaFold quality levels in different RMSD intervals on the benchmark set; E) Distribution of TMAlphaFold quality levels in different TMScore intervals on the benchmark set; F) Distribution of pLDDT values in the transmembrane regions in different TMAlphaFold quality levels on the benchmark set.

Next, we wanted to find out whether these filters really detect structures with errors. For proteins with solved 3D structures, we made a 2D projection to define their topography (the position of the TM helices in the amino acid sequence). We compared the experimentally determined or predicted topography with the topography we derived from the AF2 structures, after modelling the position of the membrane bilayer using TmAlphaFold structures. We defined modelled structures as correct where the topographies agreed (the number of TM segments was equal and all helices were covered with at least 5 amino acids). We examined the performance of different filters perform on correct and incorrect structures (Figure 1B). Approximately 80% of the proteins pass each filter when the structure is correct, while this is always lower for incorrect structures, ranging from 0-77%. TMAlphaFold uses 5 quality levels based on the filters, categorizing structures from failed to excellent. Excellent quality proteins are indeed topographically correct in 97% of the cases, while the number of errors increases in lower quality categories (Figure 1C). It is also visible that, considering the benchmark set, 40% of the proteins belong to the Excellent category, and less than 10% of the proteins have Poor or Failed categories.

For proteins for which 3D structure is available, we also calculated the Root Mean Square Deviation (RMSD) and TMScore using TMAlign^13^. When the experimental and predicted structures have a low RMSD (<1), almost 90% of the proteins have Good or Excellent quality, while the higher the RMSD, the lower the proportion of Good and Excellent structures (Figure 1D). The same is true for the TMScore (Figure 1E), where above the 0.5 threshold, 80% of the proteins have Good or Excellent quality, while below the 0.5 threshold, 59% are of Good or Excellent quality.

AF2 also provides a probability value for each predicted residue, called the predicted Local Distance Difference Test (pLDDT): the distribution of these values for TM regions shows that AF2 is more certain for higher quality predictions (Figure 1F).

### How AlphaFold2 shaped the coverage of the human transmembrane proteome

Next, we extended our analysis to the entire human TM proteome. First, we examined how the structural coverage of the TM proteome changed after AF2 was released. For this task, we used HHBlits^14^ to search for PDBTM structures where the deposited structure covered all TM helices. For each year, we only incorporated PDB^15^ structures that were available at that time. We categorized the hits based on sequence identity and created a graph to visualize the data (see Figure 2A). In the early 2000s, the number of solved structures with high sequence identity (>80%) was less than 100. Even when considering very distant homologs (<20%), about 20% of the TM proteome had a recognizable paralog. Notably, this also marked the upper limit of proteins for which 3D structure prediction could be performed using homology-recognition or threading approaches. As more research groups gained access to cryo-EM, the number of close (and distant) homologs increased, showing a promising trend. However, it would have taken decades to cover the entire TM proteome with high-quality templates. If we add AF2 structures to this graph using the TmAlphaFold quality score, the impact of AF2 becomes readily apparent, with more than half of the TM proteome falling into the Good and Excellent categories (see Figure 2A).

**Figure 2.**
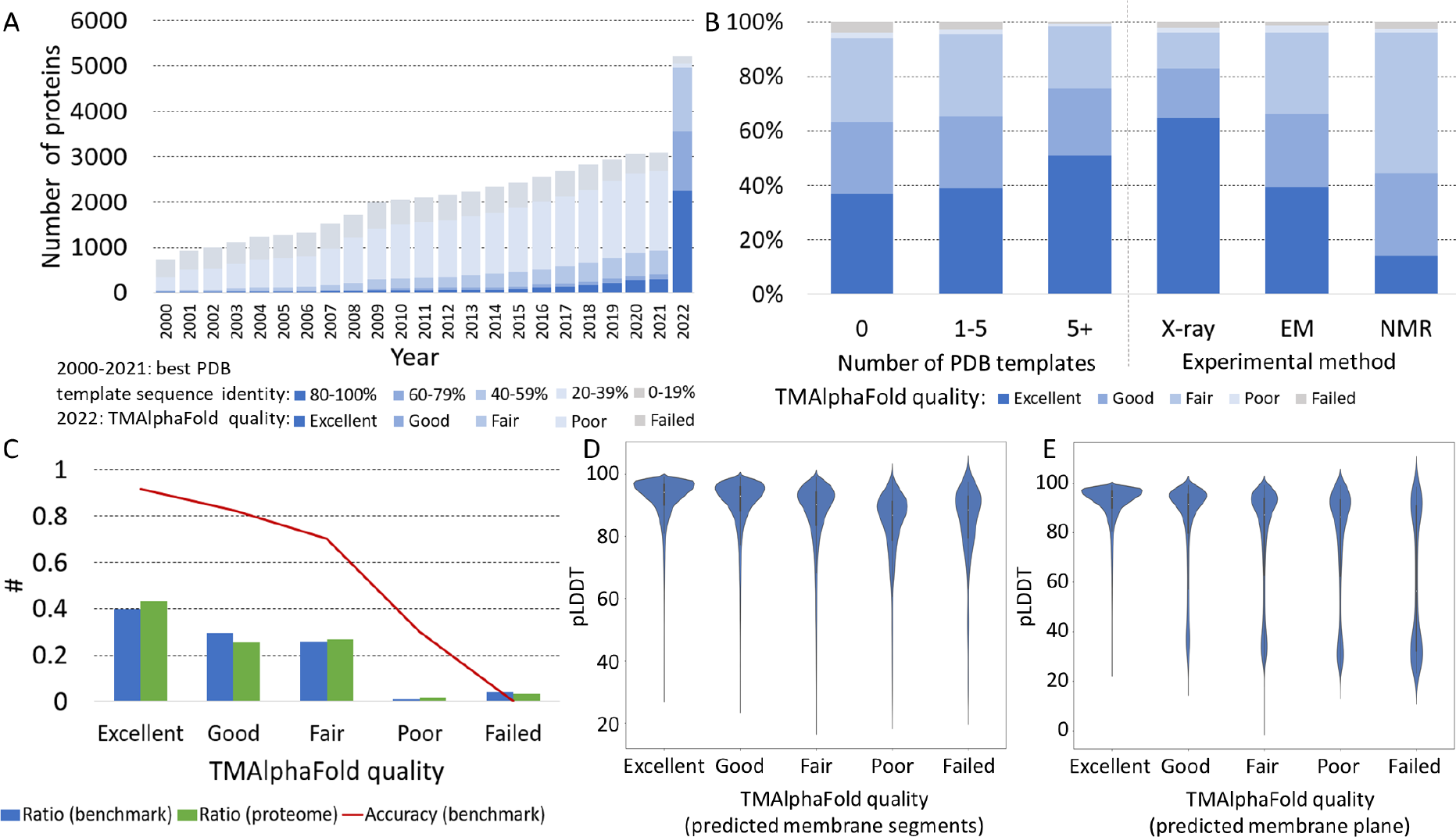
A) Coverage of PDB structures (from 2000-2021) according to their sequence identity and TMAlphaFold (2022) structures based on their quality level on the Human Transmembrane Proteome; B) Distribution of TMAlphaFold quality levels with and without homologous structures and based on experimental technique; C) Distribution the number of TMAlphaFold structures in different quality levels (benchmark set results are also displayed for comparison); D) Distribution of pLDDT values in the HTP predicted transmembrane regions in different TMAlphaFold quality levels; E) Distribution of pLDDT values in the TMAlphaFold detected transmembrane regions in different TMAlphaFold quality levels.

Although AF2 performs quite well, it is a supervised method, and this becomes evident when we check the quality of proteins for which no homologous protein was available in the PDB and for which 1-5 or more template structures were already solved (see Figure 2B). The number of excellent structures is slightly higher when there are a few homologs, but AF2’s performance is convincing when there are at least five homologs. AF2 has been trained on X-ray structures and therefore performs better on proteins with available X-ray templates than on proteins with only NMR or cryo-EM homologs in the PDB (see Figure 2B).

The distribution of proteins in different quality categories on the benchmark set and the whole proteome is almost identical; therefore, we can assume that about 40% of the whole TM proteome has a 97% accuracy (considering topography), and about 90% of the TM proteome has a 70% accuracy (see Figure 2C).

We also calculated the distribution of pLDDT values on (I) predicted TM regions (see Figure 2D) and (II) within the modeled lipid bilayer according to TMAlphaFold (see Figure 2E). In both cases, the higher the structure quality, the higher the mean pLDDT value in the membrane regions. In the case of the modeled lipid bilayer, the distribution is bimodal on all quality levels except Excellent, probably reflecting the flexible protein segments that violate the membrane. Since the lipid bilayer was not defined for AF2 training, disordered regions often cross the bilayer, resulting in incorrectly located domains.

### High quality AlphaFold2 structures provides a solid base for fast and accurate searching and clustering

Searching for distant homologs and clustering protein sets are commonly used tools to associate annotations or generate benchmark sets. In this study, we aimed to compare traditional homology searching tools with Foldseek^16^ clustering, based on AlphaFold2 structures. Accordingly, we collected protein sets from several databases, including CATH^17^, UniProt^18^, Membranome^19^, and the Transporter Classification Database^20^. We then manually discarded proteins or merged groups to create protein families with the same architecture (e.g. number of TM segments).

We first searched for homologs using HHBlits, BLAST^21^, and Foldseek (using the full AF2 structures and also using that parts of the structure that are within the TMAlphaFold predicted bilayer +-5 Ångström). In all cases, we used two approaches: one where all hits were accepted, and another where only hits with the same number of TM segments were accepted. For sequence-based approaches, we used topologies from the HTP database, while for AF2 structures, we used topologies derived from TMAlphaFold structures. The sequence and structure-based searches yielded similar results (Table 1): in both cases, the best achievable sensitivity was around 0.87, and the best specificity was close to 1. Considering more universal metrics that take into account both positive and negative cases (Matthew’s Correlation Coefficient: MCC and Balanced Accuracy: BACC), HHBlits and Foldseek provided the best results.

**Table 1:** Homology searching results using different approaches. (TP: True positive, FP: False positive, FN: False negative, TN: True negative, Sens: Sensitivity, Spec: Specificity, BACC: Balanced Accuracy, MCC: Matthew’s Correlation Coefficient. AUC: Area Under Curve (based on e-value for HHBlits, BLAST and Foldseek, and probability for NNSearch)

| Approach | TP | FP | FN | TN | Sens | Spec | BACC | MCC | AUC/prob. | AUC/seqID | AUC/TMScore |
| --- | --- | --- | --- | --- | --- | --- | --- | --- | --- | --- | --- |
| HHBlits all | 24456 | 4227 | 3424 | 73463 | 0,877 | 0,946 | 0,911 | 0,815 | 0,915 | 0,932 | x |
| HHBlits Tms covered | 23315 | 3137 | 4565 | 74553 | 0,836 | 0,960 | 0,898 | 0,810 | 0,894 | 0,905 | x |
| BLAST all | 21979 | 597 | 5901 | 77093 | 0,788 | 0,992 | 0,890 | 0,837 | 0,893 | 0,893 | x |
| BLAST Tms covered | 20316 | 73 | 7564 | 77617 | 0,729 | **0,999** | 0,894 | 0,813 | 0,859 | 0,859 | x |
| Foldseek all | 24466 | 3528 | 3414 | 74162 | 0,878 | 0,955 | 0,916 | 0,831 | 0,932 | x | 0,931 |
| Foldseek Tms covered | 20715 | 992 | 7165 | 76698 | 0,743 | 0,987 | 0,865 | 0,814 | 0,797 | x | 0,891 |
| Foldseek_mem all | 21326 | 294 | 6554 | 77396 | 0,765 | 0,996 | 0,881 | 0,831 | 0,882 | x | 0,881 |
| Foldseek_mem Tms covered | 16884 | 165 | 10996 | 77525 | 0,606 | 0,998 | 0,802 | 0,723 | 0,802 | x | 0,802 |
| NNSearch | 25218 | 4168 | 2662 | 73522 | **0.905** | 0.946 | **0.925** | **0.837** | **0.985** | x | x |

Sequential and structural search approaches have an advantage for different proteins and protein sets - for example, structural approaches are likely to fail if the structures provided are of poor quality. To overcome this problem we developed a simple neural network (NNSearch) using search results from all methods that were added as an input. To avoid possible overfitting, several neural networks were built, each time using a single protein family for testing, and all other protein families for training. NNSearch achieved the highest sensitivity, balanced accuracy, MCC (Table 1). Looking at the number of missing pairs (i.e. false negatives) with “Fair” or worse quality, Foldseek struggles to find relationships in these cases compared to HHBlits (Figure 3A). By combining the sequential and structural search modes, poor-quality structures have a lower impact on finding homologs. Similarly, the number of hits found (i.e. true positive) with “Fair” or worse quality is the lowest with Foldseek, and the highest with NNSearch (Figure 3A).

**Figure 3.**
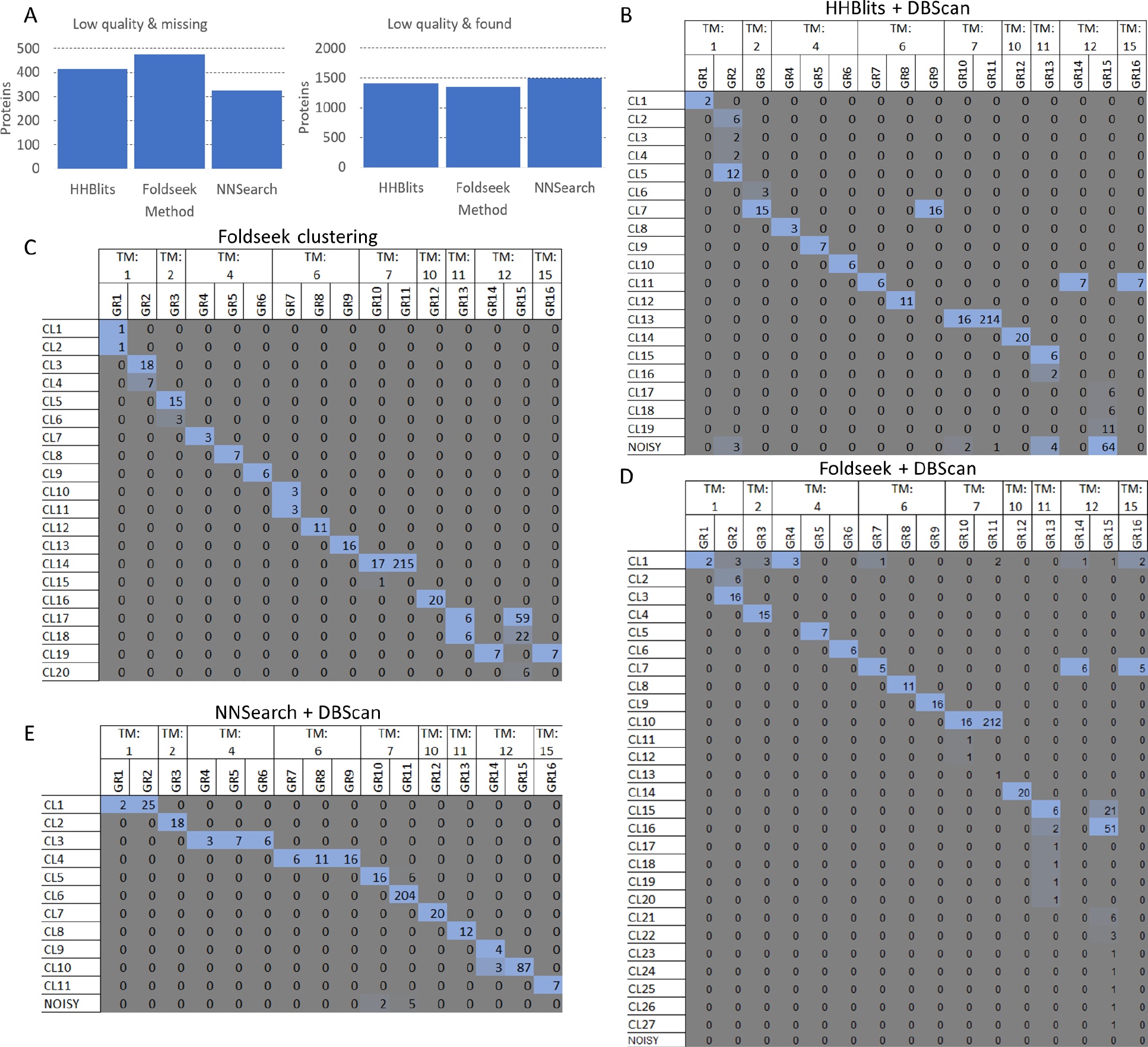
A) Left: Number of missing proteins (false negative) with “Fair” or worse quality using different approaches. Right: Number of found proteins (true positive) with “Fair” or worse quality using different approaches. B) Number of proteins in different clusters categorized by protein families (GR1: Single helix bin, GR2: Integrin, GR3: KCN_3, GR4: Eic/Glu me. Domain, GR5: Neurotim ion-channel, GR6: CACN_2, GR7: ABC_2, GR8: Aquaporin, GR9: KCN_1, GR10: Rhodopsin, GR11: Olfactory, GR12 :ATP_1, GR13: SLC_1, GR14: ABC_1, GR15: SLC_2, GR16: ABC_3), using HHBlits and DBScan. C) Number of proteins in different clusters categorized by protein families, using Foldseek clustering. D) Number of proteins in different clusters categorized by protein families, using Foldseek and DBScan. E) Number of proteins in different clusters categorized by protein families, using the Neural Network and DBScan.

Next, we used DBScan^22,23^ on HHBlits to cluster protein pairs into groups (Figure 3B). To select the best clustering result from multiple settings, we calculated the Adjusted RandIndex (ARI) and the Adjusted Mutual Information (AMI) after iterating through all reasonable parameters. Not unexpectedly for deep homology search algorithms, several families that were separated by topology in the benchmark set ended up in the same cluster, however, it cannot be taken as a serious error, considering that architectures repeating the same domain(s) but different numbers of occurences were found in these cases (e.g. ABC protein families).

We then used the built-in clustering approach of Foldseek (Figure 3C) on the full structures: its performance was superior to sequential homology search and DBScan clustering. Some protein families were split into two groups. Merging of different groups occurred mainly for ABC proteins and GPCR family members. The results can be adjusted by adding topology filters that do not allow proteins with different topologies to be in the same cluster. We also performed DBScan clustering on pairs generated by Foldseek (Figure 3D), which gave similar results to the built-in clustering method, except that it split the SLC_2 family into 4 clusters.

Finally, we performed DBScan clustering on the result of NNSearch (Figure 3E). GPCR-s are by far the largest membrane protein families and this approach successfully separated olfactory receptors and rhodopsins, but other protein families with the same number of TM segments were clustered together.

DBScan clustering on the Neural Network yielded the highest ARI (0.92) and AMI (0.90) scores, before HHBlits (ARI: 0.84; AMI: 0.82), Foldseek (ARI: 0.83; AMI: 0.79), while the FoldSeek clustering alone achieved 0.85 ARI and 0.85 AMI score.

### Areas where AlphaFold2 has limited accuracy

We also looked for groups of proteins for which AF2 gave confusing results. For this task, we used NNSearch that is based on HHBlits, BLAST, and Foldseek and clustered the human proteome using DBScan. DBScan categorized over a thousand proteins into the noisy cluster, and there is also a suspiciously large group dominated by bitopic membrane proteins. Altogether there are 197 protein clusters. For further analysis we converted the TMAlphaFold quality level to a scale of 1-5 (from failed to excellent). There are 62 protein clusters, where the mean quality is below 3.5, containing 353 proteins in total. 40 clusters (146 proteins) of these groups do not have any 4 (Good) or better quality member. Proteins belonging to these clusters belong to the most elusive part of the TM protein space, and further experimental/computational work on them are desired to better understand their function. There is an additional 31 clusters and 1763 proteins with mean quality below 4. Notably, all of these clusters have at least one representative member with 4 or better quality.

There are several other protein families with generally poor quality structures, such as members of the tumor necrosis factor receptor superfamily, of the solute carrier family, glutamate receptors and others. Looking at the GeneOntology molecular function (Figure 4A) analysis for Fair quality structures or below (using all human TM proteins as a background), terms related to kinase activity are enriched (proteins involved are likely to be disordered). For biological processes (Figure 4B), cell adhesion (typically bitopic membrane proteins), protein phosphorylation (high disordered content) and synaptic/neuronal process related terms (large proteins with multiple modules) are enriched. For the cellular compartment cell surface (many bitopic proteins) and receptor complex (multimeric proteins) terms were enriched.

**Figure 4.**
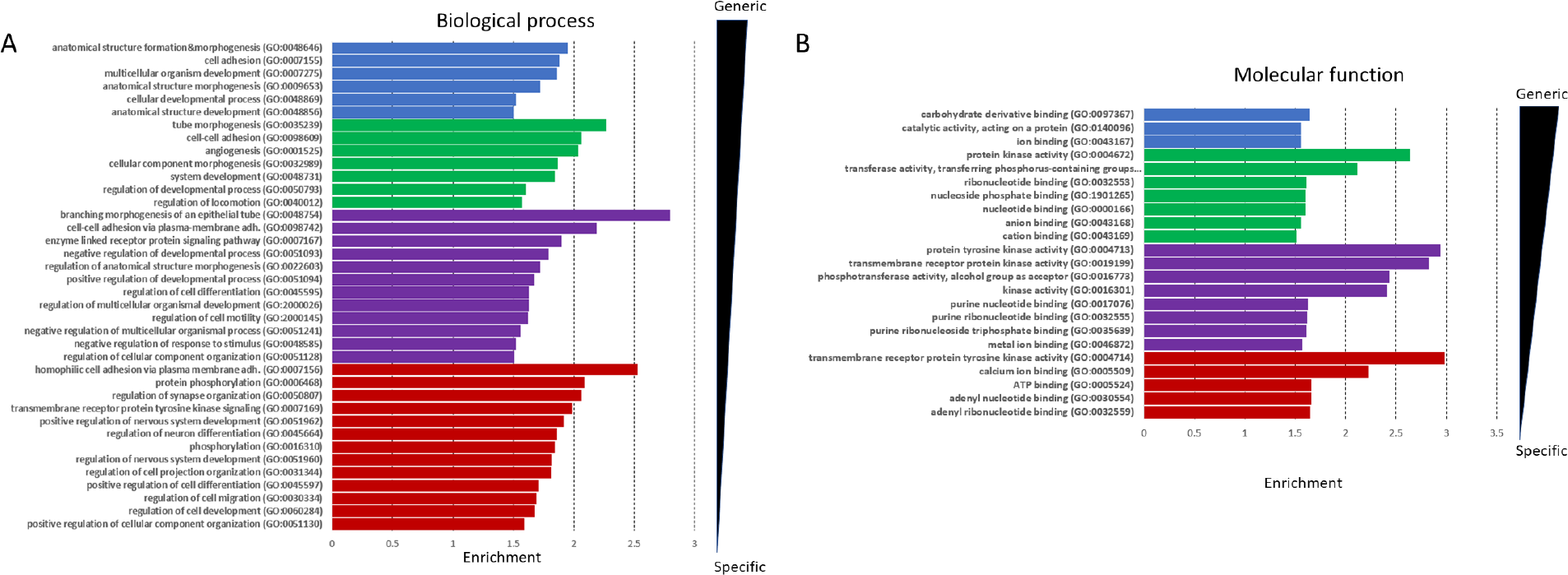
Highly significant terms are sorted based on their level in the GeneOntology tree (blue, green, purple, red: 2-5, respectively) and on fold enrichment. D: Molecular Function E: Biological process

## Discussion

In this study, we evaluated the quality of alpha-helical membrane protein structures predicted by AlphaFold2. Although AF2 has made a major contribution to the coverage of the human transmembrane proteome, there are still protein groups for which its performance seems to be poor. Scientists often analyze their protein of interest or research focus, and in these highly studied areas, AF2 performs better (as these proteins have usually been studied for decades), for example in the case of ABC transporters^26^. AF2 currently relies heavily on (X-ray structures in) the PDB, as the quality of predicted structures dropped dramatically when we considered proteins for which no template structure could be found. About 30% of the human proteome still lacks structures of convincing quality. One area where AF2 is limited is in multimeric proteins, where the correct fold is highly dependent on interactions. Although AF2 was quickly tailored by the scientific community to accept multiple protein sequences^27^, this is not yet reflected in the AlphaFold Protein Structure database^28^. Proteins with high levels of disorder also cause problems for AF2, for example despite the relatively high number of solved syndecans^29^, their quality lags behind due to the flexible regions of their ectodomains.

Defining protein clusters is also a challenging area. Multiple structure-based and guided methods have recently developed^30,31^ to improve homology searches, but all of these algorithms rely heavily on the quality of the input structures. In the case of TM proteins, we found that it is a good idea to use AF2 when the quality of the proteins is high (e.g. the TmAlphaFold database can be a good starting point), but classical sequence-based homology search will outperform structural methods when the quality is lower.

In general, AF2 seems to have three main problems when predicting membrane proteins: I) Flexible connecting loops often cross the bilayer and sometimes floating helices are also loosely placed into the membrane, especially in the case of bitopic TM proteins. II) Membrane segments are predicted outside the lipid bilayer - most often followed by a flexible segment. III) AF2 forces compact structures and folds non-TM alpha helices to the membrane plane. The following examples are not only clusters with multiple poor-quality structures but also represent the aforementioned errors.

We found several protein groups where AF2 has limited accuracy on all members. Spermatogenesis-associated proteins are expressed in spermatocytes and the retina, and some of its members are associated with retinal disease (Figure 5A). HTP predicts them as bitopic membrane proteins, but the reliability of the predictions is always below 85%. Using the AlphaFold2 structures, TMDET^24^ detects 1-5 TM segments with several failed tests, therefore the qualities of all structures are below “fair”. Inspection of the structures shows that some helical segments are present, but the low pLDDT segments connecting them often cross the membrane and place the helical segments randomly. Low pLDDT values indicate flexible disordered regions, and in this case, MemDis^25^ confirms that these proteins are highly disordered. We also found a group of cation channels with 7 TM helices, but all structures are of Poor or Fair quality (Figure 5B). Several TM segments are folded outside the lipid bilayer, and there are also non-TM regions that are incorrectly predicted to be in the lipid bilayer. In this case, numerous electron-microscopy structures are available that can be used as a template (e.g. PDB: 6MIZ). Anoctamins are anion channels that also seemed to be a challenging task for AF2 (Figure 5C). They have 8 TM segments, but AF2 tries to make the protein more compact by folding additional helical regions into the lipid bilayer.

**Figure 5.**
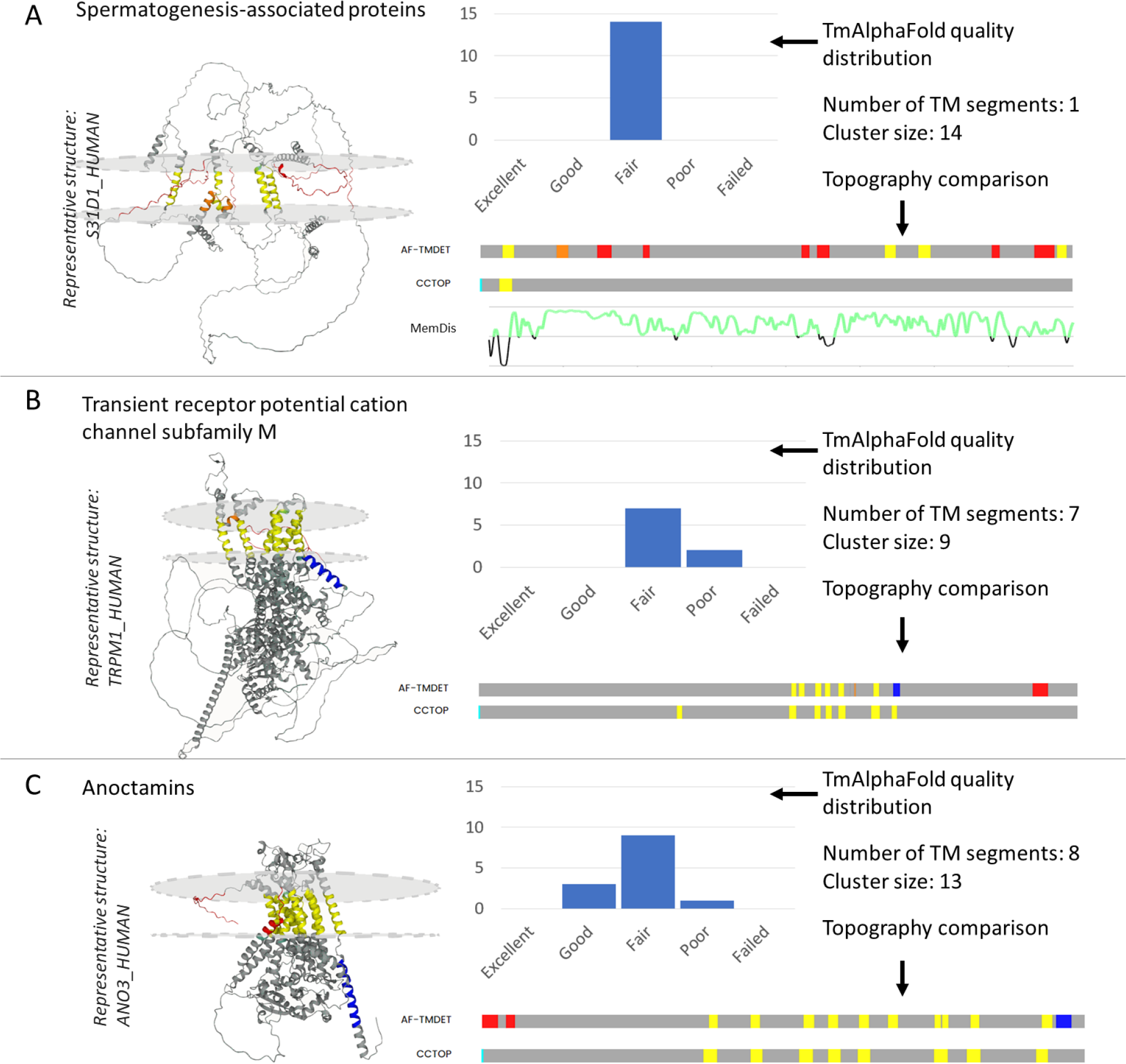
Examples of badly modelled structures from populated clusters, with a representative structure, topology comparison and TMAlphaFold quality levels. A) Spermatogenesis associated proteins. Disordered regions are also highlighted using MemDis prediction (green is disordered). B) Cation channels. C) Anoctamins.

To improve the quality of predictions, one can rely on existing PDB structures, or even on homologous AF2 predictions if their quality is high enough. To demonstrate this, we chose anoctamin-4, which has a helix and a flexible region that violates the lipid bilayer. Multiple sequence alignment to the solved anoctamin-6 structure shows that the TM regions as defined in the PDBTM database are highly conserved (Figure 6A). Using ColabFold^32^ allows users to select template mode and use one of the electron microscopy structures or the AF2 predicted anoctamin-6 as a template. Using the latter one we successfully amended the erroneous anoctamin-4 membrane domain (Figure 6B). Older modeling or threading methods, such as SwissModel^33^ or Phyre2^34^ should also not be forgotten. AF2 is trending and has a remarkable performance, but it is also a good practice to use several prediction methods and compare their performance, especially when the AF2 provided structure’s quality is suspicious. TMAlphaFold can be used to assess the quality of AF2 predicted TMP structures to pinpoint problematic proteins.

**Figure 6.**
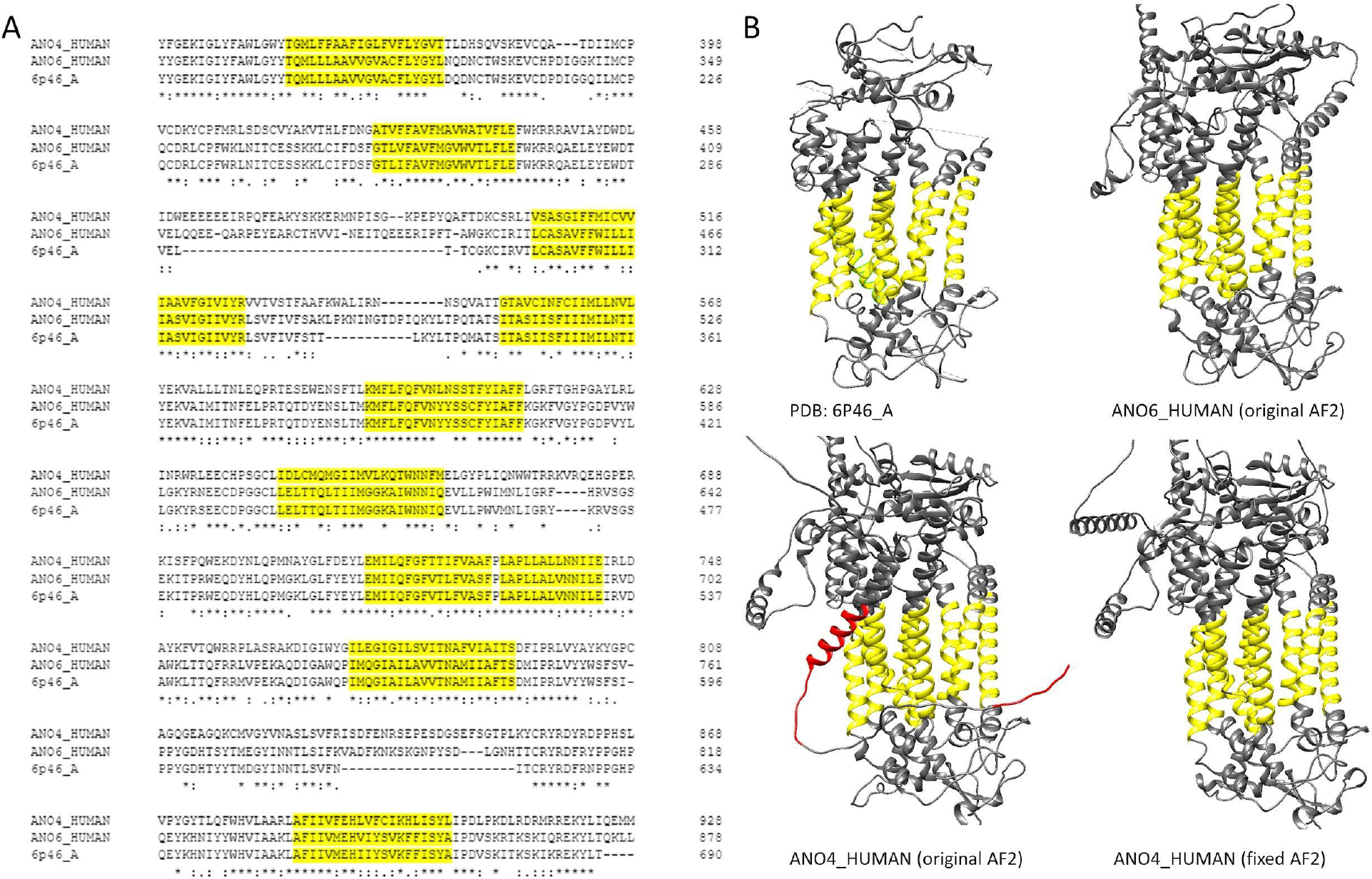
A) Multiple sequence alignment of human Anoctamin-4, Anoctamin-6 and PDB:6P46_A. B) Top: Electron-microscopy and AF2 predicted structure of Anoctamin-6, Bottom: AF2 predicted structure of Anoctamin-4; AF2 predicted structure of Anoctamin-4 using Anoctamin-6 AF2 structure as a template. Yellow regions show membrane regions as defined by TMDET. Red regions are non-TM regions placed in the plane of the lipid bilayer.

## Methods

### Performance of AlphaFold on the Human Transmembrane Proteome

First we prepared a dataset of human proteins for which reasonably high quality structural information was available from PDBTM^3^ structures or topology predictions (reliability>95% in HTP^1^). First we used BLAST^21^ with e-value 10^−5^, 40% sequence identity to collect homologous structures for proteins deposited into the HTP database. We only accepted hits that covered all TM segments. We extended this set with proteins that have a high reliability (>95%) topology prediction in the HTP. This data set was then filtered for redundancy using CD-HIT^35^, by decrementing identity (90, 70, 50, 40) and word length (5,4,3,2). The list of collected proteins can be found in Supplementary Table 1.

We used TMAlign^13^ to compare predicted and experimentally solved structures (Supplementary Table 2).

Next we compared the predicted topology from HTP and the projected topology from AF2 structures, as the membrane bilayer was defined by TMAlphaFold^12^. Correct prediction means that all TM segments overlap by at least 5 residues. The results of the TMAlphaFold filter are also visible here. (Supplementary Table 3).

### Coverage of structure on Human Transmembrane Proteins

We downloaded the full UniProt sequence of the PDBTM alpha-helical structures using SIFTS. Using these sequences we searched for homologous hits from HTP using HHBlits^14^, where all PDBTM membrane regions overlap with the HTP sequence in the alignment, with a maximum of 5 gaps allowed – however, as some topology predictions may not be accurate, the predicted transmembrane segments from the HTP were not considered (Supplementary Table 4).

TMAlphaFold quality levels (Supplementary Table 5) and the mapped structures were used to generate Figure 2A (data: Supplementary Table 6).

### Searching for homologous proteins

We used the UniProt^18^, CATH^17^, Membranome^19^ and the Transporter Classification Database^20^ to search for protein families (Supplementary Table 7). We manually processed the results to construct a dataset, using information from the above listed databases, and in addition from UniProt functional annotations, HTP topologies, PDBTM structures, TMAlphaFold and multiple sequence alignments. We discarded proteins with little or no annotation available, or merged groups where two databases had overlapping results (Supplementary Table 8).^18^

We performed several homology search strategies: BLAST, HHBlits, Foldseek^31^ (full AF2 structures and only residues in and around the bilayer detected by TMAlphaFold). We also performed the same search, but only accepted the result if the topology of the query and the hit protein matched (i.e. all TM regions overlapped by at least 5 residues).

We also developed a Neural Network that takes as input the results of all homology search algorithms (HHBlits result; HHBlits result with overlapping TM regions; BLAST result; BLAST result with overlapping TM regions; Foldseek result; Foldseek result with overlapping TM regions; Foldseek_mem result; Foldseek_mem result with overlapping TM regions), together with TmAlphaFold2 and HTP features of protein pairs (TmAlphaFold number of TM segments; TmAlphaFold quality; HTP number of TM segments; HTP reliability). We created several training and test sets - always using one protein family to test and the rest of the protein families to train (Supplementary Table 9).

We also used several clustering techniques: as protein sequence identity and structural similarity varied greatly between families, we used DBScan^22^ to cluster the results (as it is a density-based clustering method) using the similarity matrix provided by the search tools. We iterated through different epsilon values and selected the one with the highest Adjusted RandIndex and Adjusted Mutual Information. We also used Foldseek clustering on full and partial structures (with residues in and around the bilayer), using thresholds of 0.2, 0.4, 0.6, 0.8. The selection criteria were the same as for DBScan (Supplementary Table 10-13). Cluster results for the benchmark sets are shown in Supplementary Table 14.

### Searching for groups with low quality AlphaFold structures

We used the developed Neural Network (NNSearch) to define protein clusters (Supplementary Table 15). Clusters with a mean quality of less than 3.5 (transforming Failed-Excellent to 1-5 scores, Supplementary Table 16) and more than 10 proteins were further analyzed in the Discussion (Supplementary Table 17). GeneOntology analyses were performed using GOrilla^36^, by selecting genes with Fair or lower quality, and using the full human TM proteome as a background. Highly significant terms (P-value<=10^−10^) were selected and categorized according to their depth in the ontology (Supplementary Table 18).

## Acknowledgment

Ministry of Innovation and Technology of Hungary from the National Research, Development and Innovation Fund [K132522]. This project has received funding from the European Union’s Horizon 2020 research and innovation programme under the Marie Sklodowska-Curie grant agreement No 101028908.

## References

1. Dobson, L., Reményi, I. & Tusnády, G. E. The human transmembrane proteome. Biol. Direct 10, 31 (2015).

2. Bowie, J. U. Solving the membrane protein folding problem. Nature 438, 581–589 (2005).

3. Kozma, D., Simon, I. & Tusnády, G. E. PDBTM: Protein Data Bank of transmembrane proteins after 8 years. Nucleic Acids Res. 41, D524–9 (2013).

4. Varga, J. K. & Tusnády, G. E. The TMCrys server for supporting crystallization of transmembrane proteins. Bioinformatics 35, 4203–4204 (2019).

5. Dobson, L., Reményi, I. & Tusnády, G. E. CCTOP: a Consensus Constrained TOPology prediction web server. Nucleic Acids Res. 43, W408–12 (2015).

6. Kozma, D. & Tusnády, G. E. TMFoldRec: a statistical potential-based transmembrane protein fold recognition tool. BMC Bioinformatics 16, 201 (2015).

7. Hopf, T. A. et al. Three-dimensional structures of membrane proteins from genomic sequencing. Cell 149, 1607–1621 (2012).

8. Punta, M. et al. Structural genomics target selection for the New York consortium on membrane protein structure. J. Struct. Funct. Genomics 10, 255–268 (2009).

9. Varga, J., Dobson, L., Reményi, I. & Tusnády, G. E. TSTMP: target selection for structural genomics of human transmembrane proteins. Nucleic Acids Res. 45, D325–D330 (2017).

10. Jumper, J. et al. Highly accurate protein structure prediction with AlphaFold. Nature 596, 583–589 (2021).

11. Akdel, M. et al. A structural biology community assessment of AlphaFold2 applications. Nat. Struct. Mol. Biol. 29, 1056–1067 (2022).

12. Dobson, L. et al. TmAlphaFold database: membrane localization and evaluation of AlphaFold2 predicted alpha-helical transmembrane protein structures. Nucleic Acids Res. 51, D517–D522 (2023).

13. Zhang, Y. & Skolnick, J. TM-align: a protein structure alignment algorithm based on the TM-score. Nucleic Acids Res. 33, 2302–2309 (2005).

14. Remmert, M., Biegert, A., Hauser, A. & Söding, J. HHblits: lightning-fast iterative protein sequence searching by HMM-HMM alignment. Nat. Methods 9, 173–175 (2011).

15. Bittrich, S. et al. RCSB Protein Data Bank: Efficient Searching and Simultaneous Access to One Million Computed Structure Models Alongside the PDB Structures Enabled by Architectural Advances. J. Mol. Biol. 435, 167994 (2023).

16. Kim, H., Mirdita, M. & Steinegger, M. Foldcomp: a library and format for compressing and indexing large protein structure sets. Bioinformatics 39, (2023).

17. Sillitoe, I. et al. CATH: increased structural coverage of functional space. Nucleic Acids Res. 49, D266–D273 (2021).

18. UniProt Consortium. UniProt: the Universal Protein Knowledgebase in 2023. Nucleic Acids Res. 51, D523–D531 (2023).

19. Lomize, A. L., Hage, J. M. & Pogozheva, I. D. Membranome 2.0: database for proteome-wide profiling of bitopic proteins and their dimers. Bioinformatics 34, 1061–1062 (2018).

20. Saier, M. H. et al. The Transporter Classification Database (TCDB): 2021 update. Nucleic Acids Res. 49, D461–D467 (2021).

21. Altschul, S. F., Gish, W., Miller, W., Myers, E. W. & Lipman, D. J. Basic local alignment search tool. J. Mol. Biol. 215, 403–410 (1990).

22. Schubert, E., Sander, J., Ester, M., Kriegel, H. P. & Xu, X. DBSCAN Revisited, Revisited. ACM Trans. Database Syst. 42, 1–21 (2017).

23. Garreta, R. & Moncecchi, G. Learning scikit-learn: Machine Learning in Python. (Packt Publishing Ltd, 2013).

24. Tusnády, G. E., Dosztányi, Z. & Simon, I. TMDET: web server for detecting transmembrane regions of proteins by using their 3D coordinates. Bioinformatics 21, 1276–1277 (2005).

25. Dobson, L. & Tusnády, G. E. MemDis: Predicting Disordered Regions in Transmembrane Proteins. Int. J. Mol. Sci. 22, (2021).

26. Hegedűs, T., Geisler, M., Lukács, G. L. & Farkas, B. Ins and outs of AlphaFold2 transmembrane protein structure predictions. Cell. Mol. Life Sci. 79, 73 (2022).

27. Evans, R. et al. Protein complex prediction with AlphaFold-Multimer. bioRxiv (2021) doi:10.1101/2021.10.04.463034.

28. Varadi, M. et al. AlphaFold Protein Structure Database: massively expanding the structural coverage of protein-sequence space with high-accuracy models. Nucleic Acids Res. 50, D439–D444 (2022).

29. Ricard-Blum, S. & Couchman, J. R. Conformations, interactions and functions of intrinsically disordered syndecans. Biochem. Soc. Trans. (2023) doi:10.1042/BST20221085.

30. Draizen, E. J., Veretnik, S., Mura, C. & Bourne, P. E. Deep Generative Models of Protein Structure Uncover Distant Relationships Across a Continuous Fold Space. bioRxiv 2022.07.29.501943 (2023) doi:10.1101/2022.07.29.501943.

31. van Kempen, M. et al. Fast and accurate protein structure search with Foldseek. Nat. Biotechnol. (2023) doi:10.1038/s41587-023-01773-0.

32. Mirdita, M. et al. ColabFold: making protein folding accessible to all. Nat. Methods 19, 679–682 (2022).

33. Waterhouse, A. et al. SWISS-MODEL: homology modelling of protein structures and complexes. Nucleic Acids Res. 46, W296–W303 (2018).

34. Kelley, L. A., Mezulis, S., Yates, C. M., Wass, M. N. & Sternberg, M. J. E. The Phyre2 web portal for protein modeling, prediction and analysis. Nat. Protoc. 10, 845–858 (2015).

35. Fu, L., Niu, B., Zhu, Z., Wu, S. & Li, W. CD-HIT: accelerated for clustering the next-generation sequencing data. Bioinformatics 28, 3150–3152 (2012).

36. Eden, E., Navon, R., Steinfeld, I., Lipson, D. & Yakhini, Z. GOrilla: a tool for discovery and visualization of enriched GO terms in ranked gene lists. BMC Bioinformatics 10, 48 (2009).

